# Auto-antibodies against interferons are common in people living with chronic hepatitis B virus infection and associate with PegIFN non-response

**DOI:** 10.1101/2024.09.24.614789

**Authors:** DL Fink, D Etoori, R Hill, O Idilli, N Kartikapallil, O Payne, S Griffith, HF Bradford, C Mauri, PTF Kennedy, LE McCoy, MK Maini, US Gill

## Abstract

**Background and aims:** Type one (T1) and three interferons (T3IFNs) are implicated in Chronic Hepatitis B (CHB) immunopathogenesis. IFN remains the only licenced immune modulating therapy for CHB. We measured the prevalence of auto-antibodies (auto-Abs) against T1 and T3IFNs to examine the hypothesis that they impact Hepatitis B Virus (HBV) control and treatment response, as highlighted by COVID-19.

**Methods:** Our multi-centre retrospective longitudinal study accessed two CHB cohorts, auto-Ab levels and neutralisation status were measured against T1IFN and T3IFN. Associations were tested against HBV clinical parameters.

**Results:** Overall, 11.9% (33/276) of CHB patients had any detectable anti-IFN auto-Abs and 9.8% (27/276) anti-T3IFN auto-Abs, with high incidence of PegIFN*α*-induced *de novo* auto-Ab (25.7%, 9/35). However, only a minority of auto-Ab-positive sera demonstrated neutralisation *in vitro* (3/33, 9.1%). Auto-Ab-positivity correlated with higher median HBsAg levels (p=0.0024). All individuals with detectable auto-Ab were PegIFN*α* non-responders including those without auto-Ab against IFN*α* specifically.

**Conclusions:** Non-neutralising anti-IFN auto-Abs are common in CHB and associate with higher median HBsAg levels. Further prospective study of anti-cytokine auto-Ab in CHB are required to characterise association with long-term outcomes.

**Impact and implications:** HBV and PegIFN*α* individually may induce broad auto-reactivity associated with dysregulated anti-viral immune responses. Auto-Ab screening pre-PegIFN*α* and other immunotherapies may have a critical role in stratifying patient selection.

## Introduction

CHB causes major global morbidity and mortality(1). Functional cure, defined as sustained HBsAg clearance, following a period of antiviral therapy is a rare event(2). Understanding the immunobiology of naturally-resolving infection is key to developing immunotherapies for HBV-cure.HBV clearance depends on coordination of innate and adaptive immunity, including IFN signalling (3,4). Of three IFN subtypes, T1 and T3IFNs are most strongly associated with anti-viral responses through IFN-stimulated gene (ISG) regulation (5,6). The role of the diverse IFN signalling pathways in CHB immunopathogenesis remains poorly understood. Analyses of peripheral blood have suggested that IFN secretion is limited during HBV infection(7–10). Recent transcriptomic analyses of liver tissue, however, suggest that expression correlates with liver inflammation(11–13). Equivalent tissue-level analyses for T3IFN are lacking, but T3IFN is secreted by HBV-infected hepatocytes with anti-viral properties *in vitro* (14,15).

Recent studies have established the dramatic effect on mortality of acquired auto-Abs neutralising T1IFN in COVID-19 (16). This has prompted re-evaluation of auto-Abs in other infectious diseases. In CHB the impact of anti-IFN auto-Abs remains uncertain in the natural history of infection and in determining treatment outcomes. In COVID-19, auto-Abs are predominantly against T1IFNs IFN*α* and IFN*ω* but there are no studies of anti-IFN*ω* or anti-T3IFN auto-Ab in CHB.

Exogenous IFN*α* is used as a therapy in CHB, as a finite course. Polyethylene glycol-conjugated (‘pegylated’) derivates of IFN*α* (PegIFN*α*) were introduced in 2002 to improve pharmacokinetics. Prior to PegIFN*α*, 7-39% of IFN*α*-treated individuals were estimated to develop anti-IFN*α* auto-Abs which associated with non-response (17). In the PegIFN*α* era, a single study suggested that up to 47% of individuals with any prior history of IFN therapy had detectable anti-IFN*α* auto-Abs(18). The study found no association with treatment response but did not provide serology methods nor measure of auto-Ab function *in vitro*. Importantly HBsAg clearance rates post-PegIFN*α* are low but mechanisms of CHB insensitivity to IFN*α* remain uncertain(19,20). This knowledge gap is particularly relevant while IFNs continue to be included in trials aimed at HBV and hepatitis D virus cure where anti-IFN auto-Abs may compromise outcomes (17,21).

We measured prevalence and function of auto-Abs against two subtypes of T1IFN (IFN*α* and IFN*ω*) and against T3IFN (IFNλ1) in two cohorts of people living with CHB, including longitudinal samples in individuals receiving PegIFN*α*. We tested for associations between auto-Ab and clinical parameters.

## Methods

### Study population and clinical metadata

We retrospectively assayed cryopreserved sera from two London, UK CHB cohorts undergoing routine follow-up (CHB1 (n=198) from Central and North West London NHS Foundation Trust, University College London Hospitals NHS Foundation Trust and Royal Free London NHS Foundation Trust (RFL); CHB2 (n=78) from Royal London Hospital, Barts Health NHS Trust including individuals (n=36) receiving PegIFN*α* therapy). CHB1 participants were significantly older with HBeAg negative disease (Supplementary Table 1). PegIFN*α* treatment responses were defined in accordance with international guidelines(22). All clinical outcomes were obtained during routine appointments. HBV-uninfected healthy controls (HC; n=31) were recruited from university research staff. Systemic lupus erythematosus (SLE) patient samples with known anti-IFN auto-Ab were used as positive controls(23).

### Anti-IFN auto-Ab detection

Anti-IFN IgG detection was undertaken using Gyrolab microfluidic immunoassay platform as previously described with wash, capture (recombinant IFN) and detect (anti-human IgG Ab) reagents and 1:10 PBS-diluted samples(24). Results are expressed as semi-quantitative arbitrary units. Detection cut-off was defined as 2 standard-deviations below mean HC values.

### IFN signalling bioassay

To assess the ability of patient serum to block T1IFN and T3IFN pathway activation, human embryonic kidney (HEK)-293 cells were used which express luciferase, controlled by an IFN-sensitive response element (*ISRE*) sensitive to both T1IFN and T3IFN signalling(25). HEK-293-cells were cultured at 37^°^C 5% CO_2_ in DMEM media containing 10% FBS and 1% penicillin/streptomycin. Assays were performed in photometric 96 well-plates with cells seeded at a density of 2×10^5^ cells/mL. Serum was used with an end dilution of 1:10 in the presence of media (no IFN), 0.1ng/mL IFN*α*2a (hereafter IFN*α*), 0.1ng/ml IFN*ω*, or 5ng/ml IFNλ1 (hereafter IFNλ). The cells were incubated overnight, cells lysed, luciferase substrate added, and light units measured using a luminometer (BioTek Synergy H1 Multimode Reader). Samples were run in duplicate on different plates. IFN signalling activity was calculated from fold induction of samples over negative control (media only) and expressed as percentages normalised to activation by each IFN dose.

### Statistical analyses

All analyses were performed using GraphPad Prism V10.0.2 or STATA. Mann-Whitney U-test or Kruskal-Wallis test (for groups of 3 or more) were used to compare unpaired whilst Wilcoxon matched-pairs sign rank test (Wilcoxon) was used to compare paired data. Fisher’s exact test or Chi-squared test were used for contingency tables. Auto-Ab levels were log transformed as continuous dependent variables to fit a random intercepts model which accounted for clustering by individuals. Separate models were run for each auto-Ab. Given the exploratory nature of the study, univariate analysis was performed for variables. For CHB1 missing data were interpreted as missing completely at random and no substitution or imputation methods were applied. P values <0.05 were considered statistically significant.

### Ethics

Serum samples were accessed via RFL and UCL Biobank ethical Review Committee (UCL/RFL Biobank; REC reference: 11/WA/0077 or 21/WA/0388), Barts and The London NHS Trust Ethics Review Board (REC reference 10/H0715/39 or 16/LO/1699) and UCL SLE cohort study (REC reference no. 14/SC/1200).

## Results

Within the overall CHB cohort, 11.9% (33/276) subjects showed evidence of any anti-IFN auto-Ab production, with 3.6% (10/276) of these demonstrating auto-Abs against all 3 subtypes (Table 1; Figure 1C). The prevalence of auto-Abs against T1 (8.1%; 26/198) and T3IFNs (10.6%; 21/198) was higher in the CHB1 compared to CHB2 cohort (3.8%, 3/78 and 3.8%, 3/78; Table 1; Figure 1A), prior to any PegIFN*α* exposure. We noted a significant increase in the prevalence of all auto-Abs in the CHB2 cohort following exposure to PegIFN*α* therapy, notably 8.6% (3/35) developed *de novo* anti-IFN*α* auto-Abs, 14.3% (5/35) anti-IFNϖ auto-Abs, and 14.3% (5/35) anti-IFNλ auto-Abs, during longitudinal follow-up (Table 1; Figure 1B). Individuals (n=4) developing *de novo* anti-IFN*α* auto-Abs post-PegIFN*α*, sero-reverted within 6 months and one individual continued to increase auto-Ab levels post-PegIFN*α* therapy (Figure 1B). Furthermore, subjects acquiring auto-Abs against IFN*ϖ*and IFNλ remained seropositive throughout the course of follow-up (Figures 1B).

**Table 1.** Overall anti-IFN auto-Ab outcomes including PegIFN*α*-exposed participants.

|  | HC | CHB1 | CHB2 | CHB total |
| --- | --- | --- | --- | --- |
| Total | 31 | 198 | 78 | 276 |
| Anti-IFNα auto-Ab positive (n,%) | 2 (6.5) | 14 (7.1) | 6 (7.7) | 20 (7.2) |
| Median level (Gyros; 95%CI) | 11 (10-14) | 20 (18-22) | 16 (15-21) | - |
| *De novo* during PegIFN | - | - | 3 | - |
| Neutralising | 0 | 0 | 1 | 1 |
| Anti-IFNω auto-Ab positive | 1 (3.2) | 10 (5.1) | 7 (9.0) | 17 (6.2) |
| Median level | 18 (14-21) | 17 (15-19) | 16 (14-22) | - |
| *De novo* during PegIFN | - | - | 5 | - |
| Neutralising | 0 | 2 | 0 | 2 |
| Anti-IFNλ auto-Ab positive | 1 (3.2) | 20 (10.1) | 6 (7.7) | 27 (9.8) |
| Median level | 15 (10-18) | 21 (19-24) | 17 (15-21) | - |
| *De novo* during PegIFN | - | - | 5 | - |
| Neutralising | 0 | 0 | 0 | 0 |
| Any anti-T1IFN auto-Ab positive | 2 (6.5) | 16 (8.1) | 11 (14.1) | 27 (9.8) |
| *De novo* during PegIFN | - | - | 8 | - |
| Neutralising | 0 | 2 | 1 | 3 |
| Any auto-Ab | 2 (6.5) | 21 (10.6) | 12 (15.4) | 33 (11.9) |
| *De novo* during PegIFN | - | - | 8 | - |
| Neutralising | 0 | 2 | 1 | 3 |

**Figure 1.**
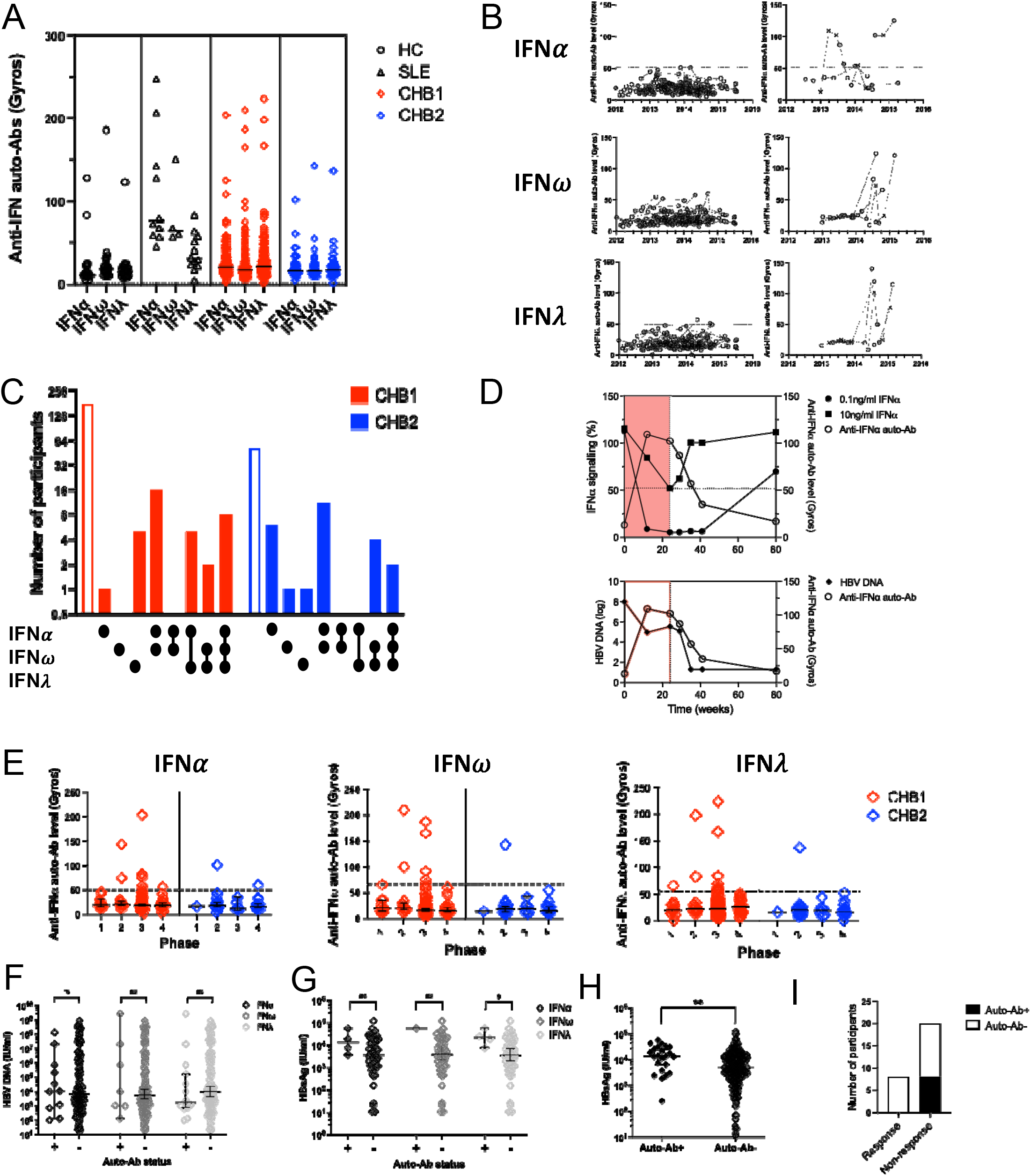
Prevalence, function and clinical associations of auto-Abs against three sub-types of IFN (1A) Scatter plot of serum auto-Ab levels against three IFN sub-types expressed as arbitrary units (Gyros). HC (healthy controls), SLE (SLE patients with known auto-Ab status), CHB1, CHB2 (n=54 without PegIFN*α* exposure). (**1B**) Line charts for serum auto-Ab levels against three IFN in longitudinal CHB2 samples 2012-2016 for individuals with no and any auto-Ab-positivity. The cut off for detection for each auto-Ab is shown by the dotted horizontal line. Current PegIFN*α* treatment is indicated by cross plots. (**1C**) Bar chart of overall participant numbers for serum auto-Ab below (hollow bars) and above the level of detection (solid bars), against three IFN sub-types with intersecting seroreactivity illustrated by upset plot connecting lines. (**1D**) IFN*α* signalling neutralisation by single participant longitudinal serum samples, with anti-IFN*α* auto-Ab levels (empty circle plots) plotted against 0.1ng/ml IFN*α* (solid circle plots) and 10ng/ml IFN*α* (square plots) signaling activation, or HBV DNA (log). Cut-off of detection for auto-Ab shown by horizontal dotted line. PegIFN*α* exposure shown by pink block prior to NUC therapy. (**1E**) Auto-Ab levels against three IFN sub-types organised by CHB phase for CHB1 and CHB2 without PegIFN*α* exposure. **(1F**) HBV DNA (IU/ml; n=36 missing data) and (1G) HBsAg levels (IU/ml; n=44 missing data) for all CHB subjects without PegIFN*α* exposure organised by auto-Ab status against three IFN sub-types. (**1H**) HBsAg levels for all CHB (with or without PegIFN*α* exposure) organised into columns with or without any detectable auto-Ab. (**1I**) Frequency of auto-Ab by PegIFN*α* response and non-response.

We tested IFN neutralisation *in vitro* and showed this was only present in one subject with anti-IFN*α* auto-Ab (1/20, 5.0%; Figure 1D), while receiving PegIFN*α*, and two individuals with anti-IFN*ϖ*auto-Ab (2/19, 10.5%), both without PegIFN*α* exposure (Table 1).

There was no difference in auto-Ab levels between CHB phases (Figure 1E) and median levels of HBV DNA in treatment-naïve individuals did not significantly differ by auto-Ab status (Figure 1F). Differences in relation to HBsAg were noted, which were significantly higher in individuals with detectable anti-IFNλ auto-Abs without PegIFN*α* exposure (22754 IU/ml, 95%CI 7815-58050) compared to individuals who had received PegIFN*α* (6370 IU/ml, 95%CI 4999-9170, p=0.0470; Figure 1G). Moreover, in patients that were treated with PegIFN*α*, HBsAg levels were greater in patients with any detectable auto-Ab (13780 IU/ml, 95%CI 6585-26067) compared to those without (5069 IU/ml, 95%CI 4234-6370, p=0.0024; Figure 1H). A significant association between anti-IFN*α* auto-Ab and HBsAg levels was noted by random intercepts modelling (coefficient 0.11, 95%CI 0.02-0.19, p=0.017; supplementary table 2). Interestingly, all individuals with any detectable anti-IFN auto-Ab demonstrated non-response to PegIFN*α* and those individuals considered responders to PegIFN*α* were seronegative for all anti-IFN auto-Abs (Figure 1I), however this association within in a small cohort did not reach statistical significance (p=0.0628).

## Discussion

In this study we demonstrate the high prevalence of auto-Abs against T1 and T3IFNs in people living with CHB which associate with impaired control of HBV replication and non-response to PegIFN*α* treatment. We noted, in our study, the incidence of developing anti-IFN*α* auto-Ab, induced by PegIFN*α* were comparable to the rates seen with then use of standard IFN prior to pegylation (7-39%)(17). Most participants with auto-Ab had reactivities against two or three IFN sub-types (21/33, 63.6%). Co-carriage of anti-IFN*α* and anti-IFN*λ*auto-Abs is as high as 51% in COVID-19 (26,27), and anti-T3IFN auto-Ab prevalence has only been measured previously in two COVID-19 studies (3.6-10.1%) (28,29) and thus prior data in CHB remains limited; our study is thus key in filling this knowledge gap.

The prevalence of neutralising auto-Ab in our study (3/276, 1.1%) remained low, but notably was still greater than reported the general population (28,30). Most binding auto-Ab did not neutralise their target antigen *in vitro*, for which there may be various reasons. Potential insensitivity of the reporter assay is a possibility, although this has been used across other studies(30) or there may be alternative non-Fab-mediated effector functions of auto-Ab. Importantly for pathogenic anti-IFNγ auto-Abs, associated with severe mycobacterial and fungal infections, Fc-mediated antagonism of T1IFN signalling and innate cell cytotoxicity have been reported(31). Thus the effector function of auto-Ab detected in our study may not be Fab-mediated to explain the apparent association with increased virus replication.

Even in the absence of specific anti-IFN*α* auto-Ab, all individuals with any auto-Ab demonstrated PegIFN*α* non-response. Carriage of any auto-Ab was associated with higher median HBsAg levels, consistent with the hypothesis that these auto-Ab may antagonise anti-viral IFN signalling leading to compromised HBV control. There is minimal structural homology between T1 and T3IFNs, which suggests that presence of auto-Abs against multiple IFN sub-types, rather than evidence of cross-reactivity, represents broad breaches of immune tolerance as seen in COVID-19. PegIFN*α* therapy is also associated with *de novo* anti-thyroid auto-Ab(32). Auto-Abs may therefore represent biomarkers reflecting an immune phenomenon predisposing to increased HBV replication that is IFN-refractory, such as increased frequency of atypical B cells which are associated with auto-Ab production and impaired anti-HBV immunity(33,34).

The major limitation of our study is its retrospective design, thus the sampling strategy was not designed to support definitive analyses for auto-Ab seroprevalence or associations with clinical outcomes. The relationship between HBsAg and auto-Ab is intriguing, however we are unable to infer direction of cause or effect. Despite longitudinal sampling, auto-Ab were frequently only detected late in follow-up which limits characterisation of the trajectory of anti-IFN auto-Ab production. Furthermore, due to the retrospective study design, we acknowledge some missing data, and although no CHB1 participants were receiving PegIFN*α* at the time of sample collection, we cannot exclude historical IFN therapy. Previous IFN treatment in some CHB1 participants may explain why overall auto-Ab profiles were comparable between the two cohorts.

In summary, our study suggests that anti-T1 and T3IFN auto-Abs are common in the UK CHB population. Our data can be used to design larger seroprevalence and immunophenotyping studies to characterise this and broader autoreactivity in CHB which may associate with treatment failure and have significant implications for future immune-mediated HBV-cure therapies. PegIFN*α*-induced neutralising auto-Abs could be risk factors for life-threatening outcomes of acute respiratory virus infection and live-virus vaccinations in the CHB population which may require enhanced monitoring.

