## Supplementary table 1 for "Auto-antibodies against interferons are common in people living with chronic hepatitis B virus infection and associate with PegIFN non-response"

|  | HC | CHB1 | CHB2 | p value |
| --- | --- | --- | --- | --- |
| Total | 31 | 198 | 78 |  |
| Age (IQR) | 36 (30-40) | 48 (41-54) | 32 (28-41) | 0.0001 |
| Sex  Male  Female  Missing | 12 (38.7)  16 (51.6)  3 (9.7) | 102 (51.5)  75 (37.9)  21 (10.6) | - | - |
| HBeAg  Positive  Negative  Missing | - | 30 (15.2)  139 (70.2)  29 (14.6) | 36 (46.1)  41 (52.6)  1 (1.3) | <0.001 |
| HBV DNA  Undetectable  Detectable  Missing | - | 29 (14.6)  134 (67.7)  35 (17.7) | 6 (7.7)  71 (91.0)  1 (1.3) | 0.04 |
| Median log HBV DNA  (IU/ml, IQR) | - | 3.17  (2.00-4.14) | 6.07  (3.77-7.54) | <0.001 |
| Median log HBsAg  (IU, IQR)  Missing | - | 3.59  (3.08-4.27)  43 (21.7) | 3.91  (3.52-4.19)  1 (1.3) | 0.1229 |
| Median ALT  (IU/L, IQR)  Missing | - | 33  (23-48)  34 (17.2) | 65  (35-129)  1 (1.3) | <0.001 |
| CHB phase  1  2  3  4  5  Unknown | - | 14 (7.1)  15 (7.6)  108 (54.5)  26 (13.1)  2 (1.0)  33 (16.7) | 8 (10.3)  28 (35.9)  14 (17.9)  27 (34.6)  0  1 (1.3) | <0.001 |
| On nucleoside treatment | - | 26 (13.1) | 35 (44.9) | <0.001 |

**Supplementary table 1. Cohort clinical characteristics**

HC= healthy controls ; CHB1= chronic hepatitis B virus infection cohort 1; CHB2= chronic hepatitis B virus cohort 2 (without PegIFNα exposure); IQR=inter-quartile range; ALT=alanine transaminase. P values generated using Kruskal-Wallis or Mann-Whitney U tests.
