## Supplementary table 2 for "Auto-antibodies against interferons are common in people living with chronic hepatitis B virus infection and associate with PegIFN non-response"

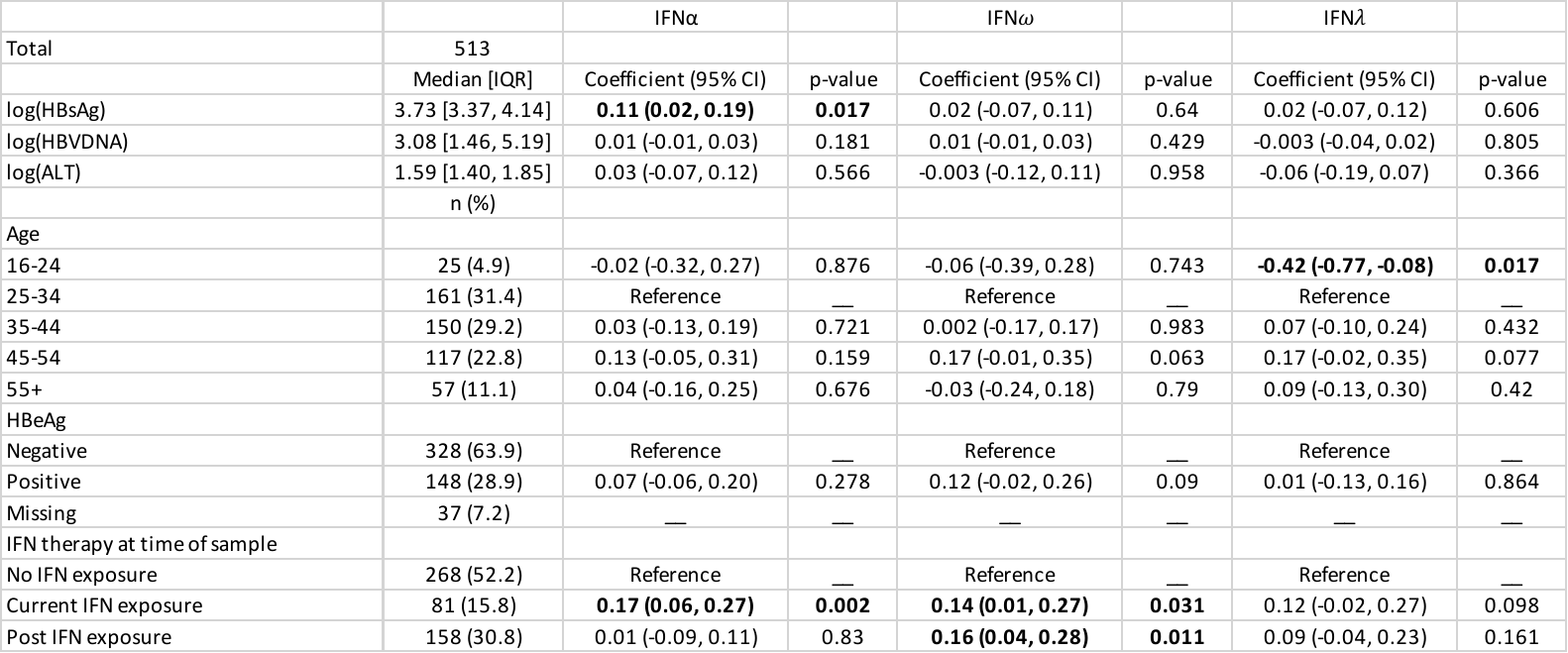

**Supplementary table 2. HBsAg and PegIFNα therapy associate with auto-Ab levels**

Random intercepts model for log transformed auto-Ab levels against IFN sub-types organised by univariate analyses for all available CHB samples (n=513). Significant interactions are highlighted in bold where p < 0.05.
